# Unpredictable soil conditions affect the prevalence of a microbial symbiosis

**DOI:** 10.1101/2023.06.07.543465

**Authors:** Trey J. Scott, Calum J. Stephenson, Sandeep Rao, David C. Queller, Joan E. Strassmann

## Abstract

The evolution of symbiotic interactions may be affected by unpredictable conditions. However, a link between prevalence of symbiosis and these conditions has not been widely demonstrated. We test for these associations using *Dictyostelium discoideum* social amoebae and their bacterial symbionts. *D. discoideum* are host to endosymbiotic bacteria from three taxa: *Paraburkholderia, Amoebophilus* and *Chlamydiae*. Three species of facultative *Paraburkholderia* symbionts are the best studied and give hosts the ability to carry food bacteria through the dispersal stage to new environments. *Amoebophilus* and *Chlamydiae* are obligate endosymbionts with no measurable impact on host fitness. We test whether the frequency of both single infections and coinfections of these symbionts are associated with the unpredictability of their soil environments by using symbiont presence-absence data from soil isolates from 21 locations across the eastern United States. We find that that *Amoebophilus* and *Chlamydiae* obligate endosymbionts and coinfections are not associated with any of our mean measures, but that unpredictable precipitation can promote or hinder symbiosis depending on the species of *Paraburkholderia* symbiont.

## Main Text

The evolution of cooperation varies with ecological unpredictability. For example, the prevalence of cooperative breeding in birds is associated with unpredictable environmental conditions [1, 2]. Cooperative breeding is thought to allow organisms to invade unpredictable environments [3] or buffer against times when conditions are harsh [4]. So far studies on the relationship between ecological unpredictability and cooperation have focused on interactions between members of the same species [1, 2, 5]. Associations between different species in a symbiosis has been suggested to have similar benefits in unpredictable environments [6–9] and may thus be associated with them. However, this association has not been tested.

We investigated whether symbiosis was associated with unpredictable conditions using the microbiome of *Dictyostelium discoideum. D. discoideum* can host three species of facultatively endosymbiotic *Paraburkholderia* bacteria. *Paraburkholderia* allow host spores to carry other species of edible bacteria and seed out food bacteria populations after dispersal at the cost of reduced spore production when edible bacteria are common [10–13]. Two of these *Paraburkholderia* species, *P. hayleyella* and *P. bonniea*, may have a longer history of host association as shown by their reduced genomes, while *P. agricolaris* may be a newer symbiont [14].

*D. discoideum* also harbors obligate endosymbionts: one from the genus *Amoebophilus* and different haplotypes from the phylum Chlamydiae. These obligate endosymbionts do not measurably affect host fitness, even when they occur as coinfections with *Paraburkholderia* [15]. We will refer to these obligate endosymbionts as *Amoebophilus* and Chlamydiae as they have not been described at the species level.

Environmental sampling has found that *Paraburkholderia* prevalence is about 25% of sampled hosts but varies by sampling location [16]. Obligate endosymbionts are found in about 40% of sampled hosts [15]. *Paraburkholderia* and *Amoebophilus* coinfections are more common than expected[15].

Host *D. discoideum* and its microbiome may be affected by unpredictable bouts of precipitation and other soil characteristics. Precipitation can drastically shift the soil environment because of the complex structure and physical properties of the soil [17]. Such shifts are known to affect the abundance of microbes in the soil [18]. When precipitation is unpredictable, it is likely to impact the availability of soil bacteria for *D. discoideum* to eat. Unpredictable precipitation may also have different effects on the soil environment depending on other soil characteristics like pH, temperature, and nutrient content. We thus suspect that unpredictable precipitation, and possibly interactions with other soil characteristics, could affect the prevalence of symbiosis in *D. discoideum*.

To test for relationships between soil characteristics and symbiont prevalence, we used presence-absence data of symbionts that were collected from 22 collection trips to 21 locations (Figure 1) across the eastern United States [15, 16]. Because some coinfections are known to be more common than expected [15], we first tested all screened hosts for non-random coinfections that may also vary with the soil environment (Figure S1, Table S1). *P. hayleyella* and *Amoebophilus* coinfections are more common than expected across our sampled sites (Figure S1). This extends prior findings that focused on a subset of locations [15]. *Amoebophilus* and *Chlamydiae* coinfections are less common than expected across our sampled sites. The rarity of *Amoebophilus* and *Chamydiae* coinfections may indicate competitive exclusion inside *D. discoideum* hosts. The association between *P. hayleyella* and *Amoebophilus* suggests that the abundance of both may be driven by the same environmental conditions.

**Figure 1:**
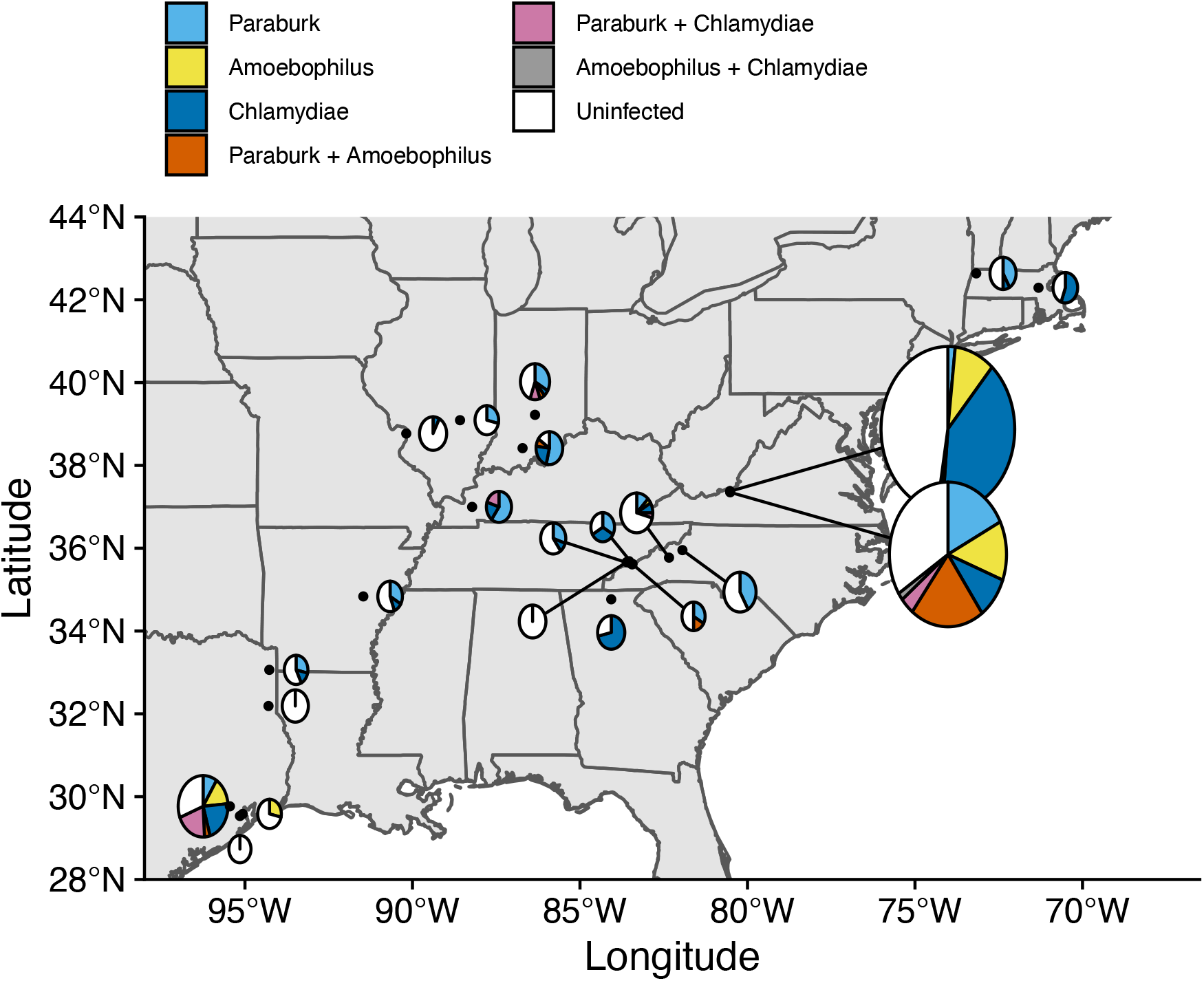
Map of *D. discoideum* sample locations and symbiont prevalence. Black points show locations. Pie charts show the frequencies of symbionts in screened hosts. Relative pie chart size indicates the number of sampled hosts at a location.

To identify associations with symbiont prevalence, we used logistic regression models (for details see SI methods).To measure precipitation unpredictability, we calculated Colwell’s P (see Table 2 in reference 20) using monthly precipitation data for each location since 1901. Colwell’s P ranges from 0 to 1, with 0 being unpredictable and 1 being perfectly predictable [19]. Along with our measure of unpredictability, we collected mean annual precipitation, soil mean annual temperature, soil carbon to nitrogen ratio, and soil pH data. We suspected that these mean soil characters that do not capture environmental unpredictability could interact with our measure of unpredictable precipitation or otherwise impact the prevalence of symbionts.

We found that the frequencies of the two *Paraburkholderia* species with reduced genomes, *P. hayleyella* and *P. bonniea*, were associated with unpredictable precipitation (Figure 2A&B; Table S2-S3). Other variables measuring mean soil characters were not associated with prevalence (though models of *P. hayleyella* prevalence showed modest support for two interactions with unpredictable precipitation; see Table S2). Interestingly, *P. hayleyella* and *P. bonniea* prevalence responded differently to unpredictable precipitation. *P. hayleyella* prevalence was higher in more unpredictable environments (log-odds = −1.066, se = 0.545; Figure 2A) while *P. bonniea* prevalence was higher in more predictable environments (log-odds = 1.202, se = 0.450; Figure 2B). For the other symbionts – *P. agricolaris*, the obligate endosymbionts, and *P. hayleyella*-*Amoebophilus* coinfections – prevalence was not associated with unpredictable precipitation or mean soil characteristics (Tables S4-S6).

**Figure 2:**
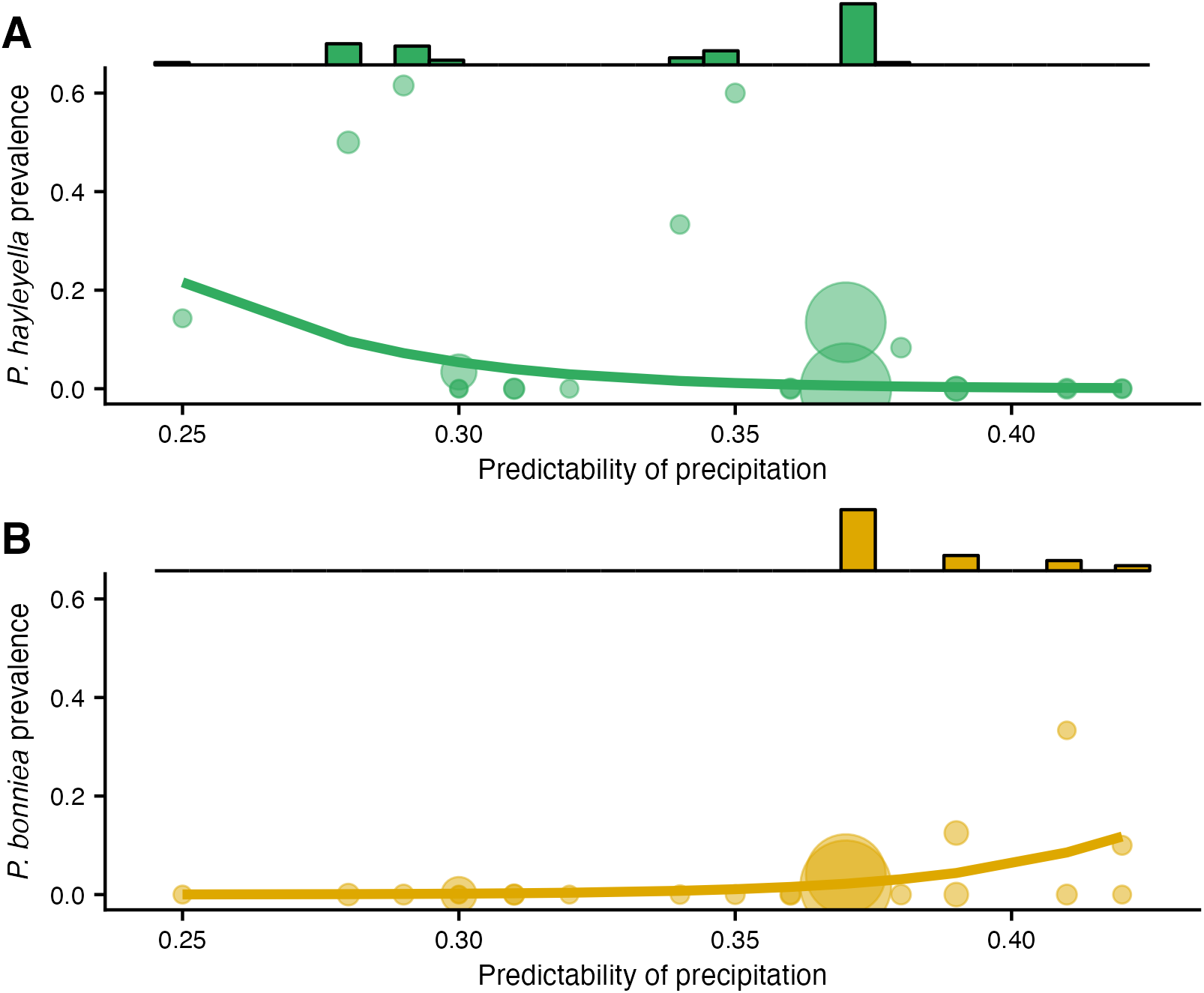
*P. hayleyella* and *P. bonniea* are differently affected by and inhabit different areas of precipitation predictability. Prevalence of *P. hayleyella* (A) and *P. bonniea* (B) for different value of predictability of precipitation along with logistic regression fits. Prevalence values are shown for each location with the size of the shape being proportional to the number of screened hosts at a site. *P. hayleyella* and *P. bonniea* inhabit different soils in terms of their precipitation predictability (histograms show number of symbionts in screened hosts at a given value of precipitation predictability; Permutation test p < 0.001).

Our finding that *P. hayleyella* prevalence increases in unpredictable conditions supports our hypothesis that symbiosis may buffer hosts during times when conditions are unpredictable. However, our finding for the relatively less common *P. bonniea* is in the opposite direction of this hypothesis.

One explanation for why unpredictability differently affects *P. hayleyella* and *bonniea* prevalence is that these sister species [10] compete and are partitioning their niches based on unpredictable precipitation. Our data show that *P. hayleyella* is found where precipitation is relatively unpredictable while *P. bonniae* is found where precipitation is more predictable (histograms in Figure 2; Permutation test: p < 0.001). Some additional support for niche partitioning comes from a previous finding that *P. hayleyella* and *P. bonniea* differ on which sugars they can metabolize [10]. One argument against niche partitioning is that the prevalence of these symbionts may not be high enough for strong competition over hosts. However, it is possible that it is competition in the soil, not in hosts, that drives niche partitioning between these symbionts.

This study demonstrates that the frequency of a microbial symbiosis can be associated with unpredictable environmental conditions. Unpredictable conditions may be an important driver of cooperation between members of the same species and between different species.

## Data Availability

Data and code is available at https://gitlab.com/treyjscott/symbiont_prevalence.

## Acknowledgements

We thank the members of the Strassmann-Queller lab along with Carlos Botero, Fred Inglis, and Jonathan Losos for feedback on this project. This material is based upon work supported by the National Science Foundation under grant numbers IOS 1656756, DEB 1753743, and DEB 2237266.

## Supplemental Materials

### Supplemental Methods

#### Data Acquisition and Processing

To measure the frequency of symbiosis, we used data from prior environmental sampling [1, 2]. The first study [1] tested *D. discoideum* isolates from 21 locations (one location was sampled two separate times) for the presence of the three species of *Paraburkholderia* symbionts [3] using *Paraburkholderia* specific 16S sequencing. The second study [2] tested a similar set of *D. discoideum* isolates for *Amoebophilus* and *Chlamydiae*, but also included samples from a few additional countries. For this study, we focused only on the United States samples because sites from other countries were not well sampled and could skew the results. We used these data to construct a presence-absence variable for each *D. discoideum* clones for whether they were infected with any of the three species of *Paraburkholderia*, or *Amoebophilus*, or *Chlamydiae*.

To investigate the role of environmental predictability on the *Dictyostelium*- *Paraburkholderia* symbiosis, we acquired data on long-term precipitation, soil pH, soil organic carbon, nitrogen, and temperature for each sample location from online databases. These variables are known to affect the abundance of bacteria in the soil [4]. For each location, we collected monthly precipitation data from 1901 to 2020 from the climate research unit database version 4.05 [5]. To measure the predictability of precipitation across these monthly measures, we calculated Colwell’s P [6] using the *Colwell’s* function in the hydrostats package [7] with 12 bins corresponding to months and with log-transformed precipitation measures as in Table 2 in Colwell [6]. Colwell’s P ranges from completely unpredictable (0) to completely predictable (1). We tested two P measures meant to capture long-term and recent predictability: (1) calculated with precipitation data from 1901 to the year that a sample was collected and (2) calculated from precipitation data from 5 years before the sample was taken. These measures were largely similar and did not change any of our results, so we include only the long-term measure in the main text.

We collected soil pH, nitrogen, and organic carbon data from the SoilGrids database version 2.0 [8]. SoilGrids are soil predictions based on empirical soil measurements and are generated at 250-meter scales. We collected soil temperature variables from Lembrechts et al. [9]. Temperature data were generated by predicting deviations of soil temperatures from air temperatures at 0 to 5 cm and 5-15 cm depths. We used 0-5 cm depths for soilGrids and soil temperature data because *D. discoideum* typically resides in the top layers of soil.

#### Statistical methods

To test for associations between different symbiont species across locations, we used mixed effect logistic regression from the *lme4* package [10] in R version 4.1.2 [11]. To account for multiple observations at a location, we used location as a random effect. We treated the location that was sampled twice (Mountain Lake Biological Station) as two separate locations because soil samples were taken from different areas within Mountain Lake Biological Station and because samples were collected 14 years apart.

As a follow up to our logistic regression results across locations, we tested whether coinfections involving different *Paraburkholderia* species were random in specific locations using Fisher’s exact tests (Table S7). To perform Fisher’s exact tests, we constructed a 2×2 contingency table for each sampling location in which at least 2 of the investigated 3 symbionts were present. To correct for multiple comparisons, we adjusted p-values using Benjamini-Hochberg’s correction.

To test for associations between soil characters and prevalence of individual symbionts or coinfections, we fit a set of mixed effect logistic regression models using lme4 [10] as above. We tested models that were derived from a full model that included the mean annual temperature (MAT), carbon to nitrogen ratio (C/N), mean annual precipitation (MAP), precipitation predictability (Collwell’s P), soil pH, and first order interactions between these variables. To reduce the risk of overfitting, we only compare models with three or fewer total predictors. To identify top models among the set derived from the full model, we used AICc values [12] and examined effect sizes of model estimates. We identify uninformative models if the model does not differ from an intercept only (null) model in terms of AICc. We identify informative models if the model AICc is less than the null model by 2 or more. For multiple models that fit better than the null model, we examined models within two AICc units of the best fitting model. We identified important effects by looking for 95% confidence intervals that differed from 0. To test for spatial autocorrelation in our models, we performed a Moran’s I test on simulated residuals using the *DHARMa* package in R [13]. All models were free of spatial autocorrelation.

To test whether *P. hayleyella* and *P. bonniea* inhabit soils with different precipitation uncertainties, we used a permutation tests. We randomly shuffled host infection status (infected with *P. hayleyella* or infected with *P. bonniea*) from hosts infected by either of these species without replacement and calculated sample statistics for values of precipitation predictability across 10,000 samples. As sample statistics, we investigated the differences between both the means and medians of the two species after permutation. Both mean and median difference statistics gave equivalent results. We report the median difference p-value in the main text.

## Supplemental figures

**Figure S1:**
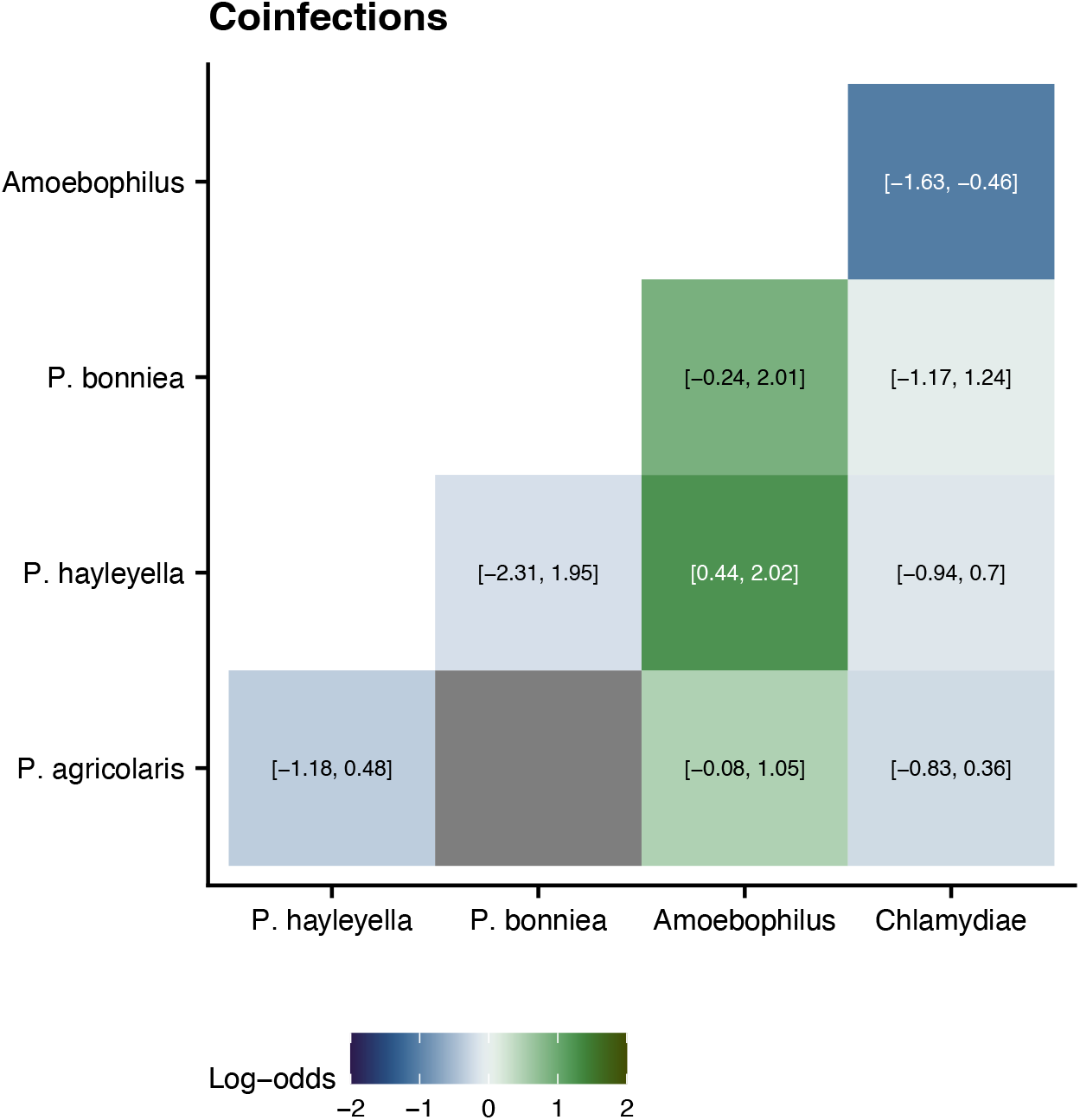
Patterns of coinfection between *D. discoideum* endosymbionts. Squares are colored according to their log-odds from logistic regression models. 95% confidence intervals are given inside squares. *P. agricolaris* and *P. bonniea* are never found together resulting in no variation for logistic regression. Fisher tests of *Paraburkholderia* coinfections can be found in the Table S7.

**Supplemental Tables (available on bioRxiv as excel file)**

**Table_S1:** Models of *P. agricolaris* prevalence

**Table_S2:** Models of *P. hayleyella* prevalence

**Table_S3:** Models of *P. bonniea* prevalence

**Table_S4:** Models of *Amoebophilus* prevalence

**Table_S5:** Models of Chlamydiae prevalence

**Table_S6:** Models of Amoebophilus-P. hayleyella coinfections

**Table_S7:** Fisher tests of *Paraburkholderia* coinfections

